# GUANinE v1.1 Reveals Complementarity of Supervised and Genomic Language Models

**DOI:** 10.64898/2025.12.06.692772

**Authors:** eyes s. robson, Nilah M. Ioannidis

## Abstract

**Background:** There has been much debate about the benefits of supervised versus unsupervised learning on genomes. Answering the question of “which is better?” requires the development of comprehensive benchmarks spanning representative functional and evolutionary tasks. Importantly, such benchmarks need large sample sizes to enable well-powered ranking of models under equivalent evaluation, such as L2-regularized probing (linear evaluation).

**Results:** Having developed such an assessment (`GUANinE` v1.1), we can conclude that the answer is both: each paradigm outperforms on certain tasks and offers key advantages over the other. Supervised sequence-to-function models excel at annotating functional states characterized by chromatin accessibility, histone marks, or CTCF binding, while unsupervised language models outperform on evolutionary conservation and related tasks without being limited to data-rich organisms. Our hundreds of new evaluations since v1.0 provide evidence for a direct tradeoff between input context size and model parameter count when on a fixed compute budget, which we depict with novel metrics like parameters/base pair. We also describe two new large-scale variant interpretation tasks: cadd-snv measuring proxy deleteriousness, and clinvar-snv measuring clinical pathogenicity. We find that conservation scores, and by extension, language models, dominate deleteriousness prediction, but successfully translating deleteriousness to pathogenicity remains challenging.

**Conclusions:** `GUANinE` v1.1 is a large-scale and thorough evaluation of pretrained genomic models. We identify uniquely performant models across tasks, and we conclude by suggesting hybrid-supervised language models may define the next era of genomic sequence modeling.

## Background

Benchmarks have a long history in computing and statistical learning—from high-performance computing [1] to computer vision [2]—and can systematically evaluate model performance across a defined, accessible suite of model assessments with defined metrics spanning multiple field-encompassing tasks.

Designing a benchmark involves careful consideration of the choice of tasks, data provenance, task breadth, and *construct validity*, which measures the claim that a singular task represents an underlying research concept. For example, a benchmark task designed to evaluate chromatin accessibility based on a single ATAC-seq experiment in a single cell line may yield starkly different results from an alternative accessibility task constructed with data from another cell line or different experiment type, e.g. DNase-seq or FAIRE-seq.

Our previously described benchmark, `GUANinE` v1.0 [3], included tasks deliberately designed to model tissue-agnostic signal aggregated across many experimental cell lines, rather than relying on tissue-specific measurements. This layer of abstraction gave `GUANinE` independence from any one experiment or cell type, with the intent to increase its staying power and long-term relevance. Several other benchmarks have also been developed in the genomics field [4–7], each with its own definitions of pre-processing, standardization, and task selection. Additionally, substantial non-dataset benchmarking has been undertaken in the field, both during the release of a new model [8–10] and stand-alone or competition-style research [11–15].

A unique feature of `GUANinE` v1.0 was its statistical precision achieved through unrivaled sample sizes across tasks. This means a model could be evaluated in a controlled environment without the need to fine-tune on small sample sizes. Eliminating this confounding factor removes the advantage of less-complex models due to statistical efficiency – a well-studied statistical phenomenon [16–18]. As a result, `GUANinE` more closely resembles the asymptotic suitability of pretrained models to tasks, subject only to construct validity.

Specifically, `GUANinE` v1.0 created multiple region-level tasks split across three different domains: regulatory element annotations [19], multiple sequence alignment conservation scores [20], and synthetic yeast promoter modeling [21]. Each domain had at least two tasks – this intentional redundancy ensured robustness of results, given the then-prototypical nature of genomic benchmarks. These tasks included:

1. `dnase-propensity` (Regulatory, hg38) – tissue-agnostic chromatin accessibility task derived from DNase hypersensitivity
2. `ccre-propensity` (Regulatory, hg38) – tissue-agnostic candidate cis-regulatory element inference, based on accessible histone marks
3. `conservation30` (Evolutionary, hg38) – mammalian conservation annotation
4. `conservation100` (Evolutionary, hg38) – vertebrate conservation annotation
5. `gpra-c` (Promoter Expression, yeast) – gigantic dual-reporter assay
6. `gpra-d` (Promopter Expression, yeast) – as `gpra-c`, but in a different culture environment

A more detailed description of each task and previously evaluated models is provided in Supplemental Background Section B.

In this work, we introduce version 1.1 of the `GUANinE` benchmark, where we evaluate dozens of new models on the existing tasks of the `GUANinE` benchmark, and we create two new *variant* benchmark tasks before reporting novel results on them.

## Results

### Newly evaluated models

For brevity, we leave concrete description of each model family and model-specific details to Methods. We cluster these by source citation and present them in roughly the same order as the results. Of course, these models are not a definitive list of sequence and sequence-to-function models in the field, and we also attempted to preclude models obsolesced by their original developers (e.g. NT-v1 models). The newly reported models in the `GUANinE` v1.1 benchmark are:

1. Two ChromBPNet models, Fold 0 and Fold 3, trained on bp-resolution ATAC-seq data from the GM12878 cell lines from Pampari et al. [22]
2. ChromBPNet training signal track ENCFF180XQC Pampari et al. [22]
3. DNABERT-2 genomic language model from Zhou et al. [7]
4. PhyloGPN phylogenetic-loss gLM from Albors et al. [23]
5. All four Nucleotide Transformer v2 (NT-v2) gLMs from Dalla-Torre et al. [24]
6. Six of the GENA-LM family of gLMs, plus one version fine-tuned on Enformer’s training data, from Fishman et al. [25]
7. The contrastive dna2vec of ‘Embed-Search-Align’ from Holur et al. [26]
8. The supervised CNN Sei from Chen et al. [27]
9. Two Mistral-DNA and two ModernBERT models from Mourad and Batut [28].
10. Two of the Caduceus genomic language models, PS and PH, from Schiff et al. [29]
11. Two of the Evo2 gLMs, including a second evaluation of Evo2 7b at a higher-context size, from Brixi et al. [30]
12. Two of the gLMs from the AIDO group, as presented by Ellington et al. [31]

Additionally, as part of the newly released `cadd-snv` and `clinvar-snv` tasks, we also include a handful of variant- and splicing-specific models in our evaluations, such as:

1. CADD v1.7, an annotation-supervised deleteriousness model trained with simulation-augmented data from Schubach et al. [8]
2. Pangolin, an RNA-splicing model developed by Zeng and Li [32]
3. GPN-MSA, a conservation-centric MSA-to-function model from [33]
4. AlphaGenome, a supervised sequence-to-function model, not yet released, but described by Avsec et al. [10]

### Newly constructed tasks

The new tasks are described in detail in Methods. Briefly, the `cadd-snv` task requires models to separate frequent or evolutionarily inferred variants from ultra-rare variation – a challenging task that proxies deleteriousness. We note that the original authors of CADD and varCADD caution against over-optimizing deleteriousness prediction [8, 34], as doing so can detract from pathogenicity generalization (e.g. to `clinvar-snv` ), but we posit that the population-derived varCADD dataset is still an incredibly useful dataset for evaluating models.

The second new task, clinvar-snv, is the smallest task yet in `GUANinE` , featuring just fewer than 12,000 variants, though its nearly 6,000 pathogenic variants is still much larger than the several hundred used by a comparable benchmark [36]. This serves to further ground evaluated models, presuming they can be extended to zero-shot regimes. Note that as described in Figure 1, the underlying *construct validity* of `clinvar-snv` is <u>not</u> general pathogenicity, rather, it is simply a well-controlled subset of the splicing-rich ClinVar database used to validate model performance. We present the results of zero- and few-shot modeling on the two new tasks after our primary results in Table 3.

**Fig. 1:**
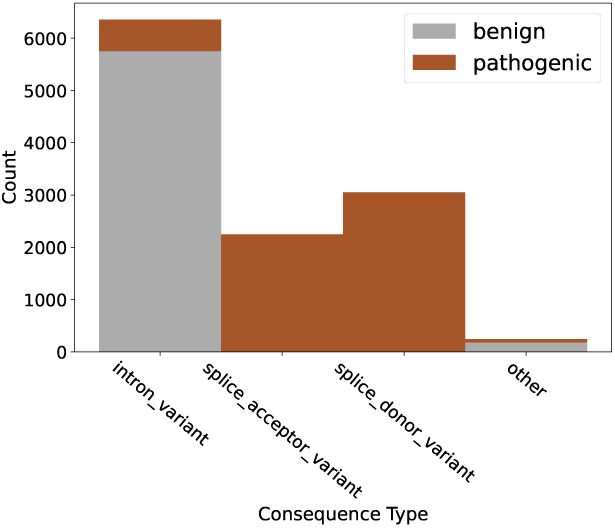
Variant consequence imbalances dominate the signal of `clinvar-snv`, which is representative of ClinVar.

### Model performance on existing tasks

The abbreviated summary of our results, which we will discuss in detail, is presented in Table 2. The full tabulation of findings, which are not as deeply discussed, is presented in Supplemental Table A1. This secondary table includes additional models we may refer to in passing, along with subtask score breakouts for the `ccre-propensity` task.

**Table 1:** Summary statistics of new variant tasks. `cadd-snv` is a large-scale dataset where models are asked to distinguish between proxy deleterious variants, singletons in gnomAD v3, and proxy benign variants, simulated evolutionarily or frequently observed in gnomAD v3 (referred to as `hfs` [34]). `clinvar-snv` asks a model to distinguish between benign variants and pathogenic variants; it predominantly consists of benign intronic and pathogenic splice site variants. While this distribution is imbalanced, it is representative of noncoding variants in ClinVar.

| Task | Proxy Deleteriousness<br>**cadd-snv** | Pathogenic Effects<br>**clinvar-snv** |
| --- | --- | --- |
| Train set | 76 M | — |
| 1% Train set | 760 K | — |
| Dev set | 1.03 M | 9.2 K |
| Test set | 1.04 M | 12.0 K |
| Split | Dev21/Test22 | Dev21/Test22 |
| Genome | hg38 | hg38 |
| Target | Boolean | Boolean |
| Output range | 0 - 1 | 0 - 1 |
| Source | Nazaretyan et al. [34] | Landrum et al. [35] |

**Table 2:**
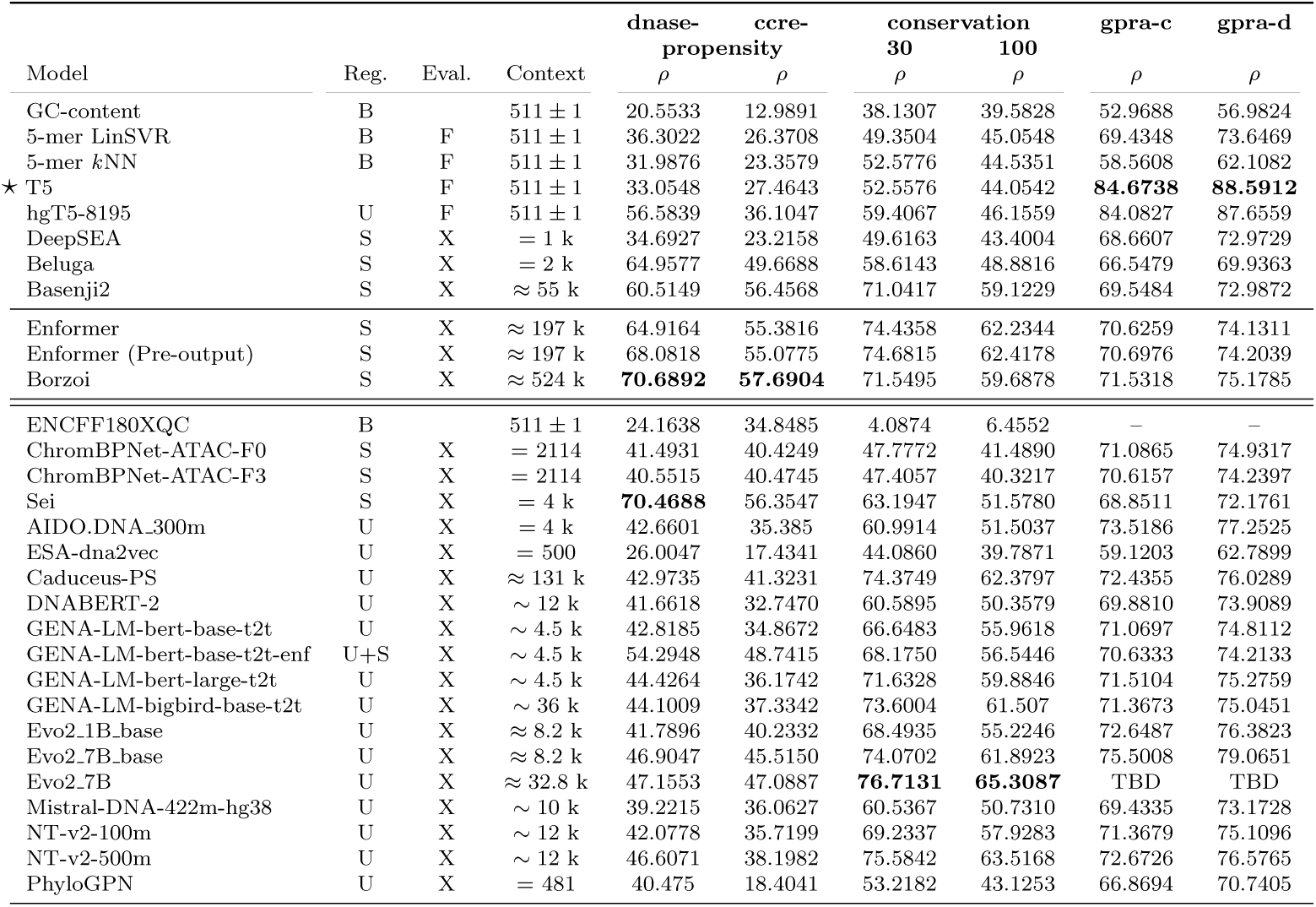
Abbreviated results on existing tasks. Models are grouped first by manuscript reported, with models below the double divider being newly reported here. Models are secondarily grouped by pre-training regime and alphabetically by family name. The best reported score, along with any close scores, are **bolded** per task. The * T5 is a fully supervised neural baseline, and the hgT5-8195 is a higher-context hg38-pretrained version of it [3]. **Tables Key:** Regimes: B → non-neural Baseline | M → MSA-based method | S → Supervised pre-training | U → Unsupervised pre-training Evaluation: F → Fully fine-tuned | X → 1% evaluation | “ ” → zero-shot Context: = → exactly | ≈ → rounded to | ∼ → on the order of

Due to the diversity of evaluated methods on new tasks, including models not suitable for region-level inference, we present their results separately in Table 3.

**Table 3:** Results on newly constructed tasks released in `GUANinE` v1.1. Note that `clinvar-snv` has no training set, so if models are not zero-shot, they are calibrated to the development set as described in Methods. The best reported score, along with any close scores, are **bolded** per task. **Tables Key:** Regimes: B → non-neural Baseline | M → MSA-based method | S → Supervised pre-training | U → Unsupervised pre-training Evaluation: F → Fully fine-tuned | X → 1% evaluation | “ ” → zero-shot Context: = → exactly | ≈ → rounded to | ∼ → on the order of

| cadd-snv |  |  |  |  | clinvar-snv |  |  |  |
| --- | --- | --- | --- | --- | --- | --- | --- | --- |
| Model | Reg. | Eval. | Context | $\rho$ | Model | Reg. | Context | $\rho$ |
| GC-content | B | | 511 ± 1 | 1.4665 | GC-content | B | 511 ± 1 | 11.5278 |
| Naïve Conseq. Type | B | X | – | TBD | Naïve Conseq. Type | B | – | 76.7449 |
| GPN-MSA | M |  |  | **29.1221** | phyloP-30way | M |  | 66.0851 |
| GPN-MSA (Position) | M |  |  | 26.6472 | GPN-MSA | M |  | 80.7211 |
| phyloP-100way | M |  |  | 23.2761 | phyloP-100way | M |  | 81.5065 |
| phyloP-30way | M |  |  | 28.7228 | PhyloGPN | U |  | 67.5918 |
| CADD v1.7 | S | | – | 18.1545 | AlphaGenome-K562 | S | ≈ 1.05 M | 93.1695 |
| CADD v1.7 (Position) | S |  | – | 19.6639 | CADD v1.7 (Position) | S | – | 83.6767 |
| ChromBPNet-ATAC-F0 | S | X | 2114 | 17.2853 | CADD v1.7 | S | – | 84.6379 |
| ChromBPNet-ATAC-F3 | S | X | 2114 | 16.7586 | Pangolin | S | ≈ 10 k | 85.6332 |
| Enformer | S | X | ≈ 197 k | 17.0984 | Enformer-Aggr. | S | ≈ 197 k | TBD |
| Sei | S | X | $=$ 4 k | 19.9110 | Sei-Aggr. | S | $=$ 4k | 50.8500 |
| Evo2.1B_base | U | X | ≈ 8.2 k | TBD | Sei-Aggr. + Pangolin | S+S | ≈ 10 k | **95.1299** |
| Evo2.7B_base | U | X | ≈ 8.2 k | TBD | | | | |
| NT-v2-50m | U | X | ∼ 12 k | 21.8891 | | | | |
| NT-v2-100m | U | X | ∼ 12 k | 23.8862 | | | | |
| NT-v2-250m | U | X | ∼ 12 k | 25.4037 | | | | |
| NT-v2-500m | U | X | ∼ 12 k | 26.4742 | | | | |
| NT-v2-500m + Sei (RF) | U+S | X | ∼ 12 k | 26.9037 | | | | |
| PhyloGPN | U | | $=$ 481 | 9.8049 | | | | |
| PhyloGPN | U | X | $=$ 481 | 23.5960 | | | | |

(a) Proxy deleteriousness is dominated by conservation scores, in particular the neural GPN-MSA and phyloP-30way. Those same conservation scores underperform on variants in **clinvar-snv** – showcasing the strategic *underfitting* to deleteriousness the CADD developers rely on for pathogenicity performance.

### Comparison of supervised and unsupervised models

First and foremost, we highlight that supervised methods like Enformer, Borzoi, and Sei excel at annotating function, as exemplified in the `dnase-prop` and `ccre-prop` results of Table 2. Specifically, consider the visible performance gap between supervised and gLM `dnase-prop` scores in Figure 2 (a). This is due in part to the data-richness of supervised element annotations in humans, which makes up the bulk of our evaluation sets. Given the relatively minor divergence of ChromBPNet-ATAC Fold 0, which sees chromosomes 21 and 22 during training, and ChromBPNet-ATAC Fold 3, which does not, we remark that there does not appear to be a significant ‘test set leakage’ effect for the supervised models being evaluated, possibly due to the robustified, cell-type agnostic nature of `GUANinE` ’s functional tasks.

**Fig. 2:**
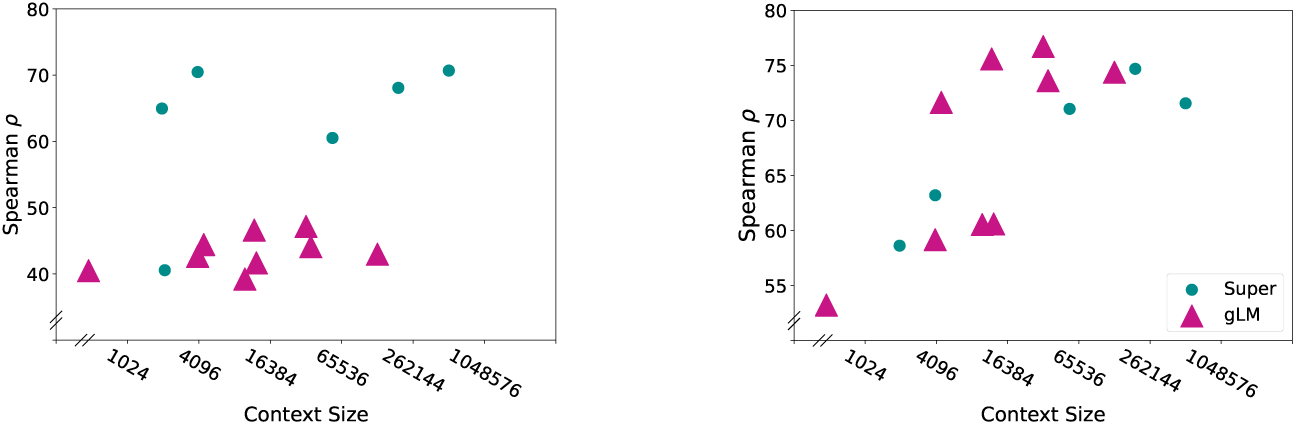
Scatterplots of task performance as a function of context size and model type. Context size, while helpful, is only weakly predictive for both functional and conservation task performance compared to other factors.

However, it is clear that a primary advantage of language models over supervised approaches is gLMs’ ability to model sequence conservation, as seen by the performance of Nucleotide Transformer v2, Evo2, and GENA-LM models on `cons30` and `cons100` in Table 2. While the gap is smaller on the cons30 task shown in Figure 2 (b), it’s worth recognizing that these gLMs have been given objectively less supervision and are likely to further outperform if fine-tuned [3, 37]. While hgT5-8195 is the only example to have been fine-tuned on conservation, we point to the functional dominance in Table 2 of fine-tuned-gLMs like hgT5-8195 and GENA-LM-bert-base-t2t-enformer over others in their respective categories.

We additionally emphasize, while not explored in `GUANinE` v1.1, the performance of multi-species gLMs on `dnase-prop` and `ccre-prop` suggest that gLMs could partially impute annotations for data-poor organisms without the need for additional supervision.

### Scaling laws apply to gLMs

On the topic of scaling laws [38], we observe that larger models tend to improve downstream model performance, while likely improving the performance-to-training-budget ratio as well [39]. We observe this in Figure 3, and note that different architectures and pretraining corpora can start a model family at different locations on the scaling law frontier.

**Fig. 3:**
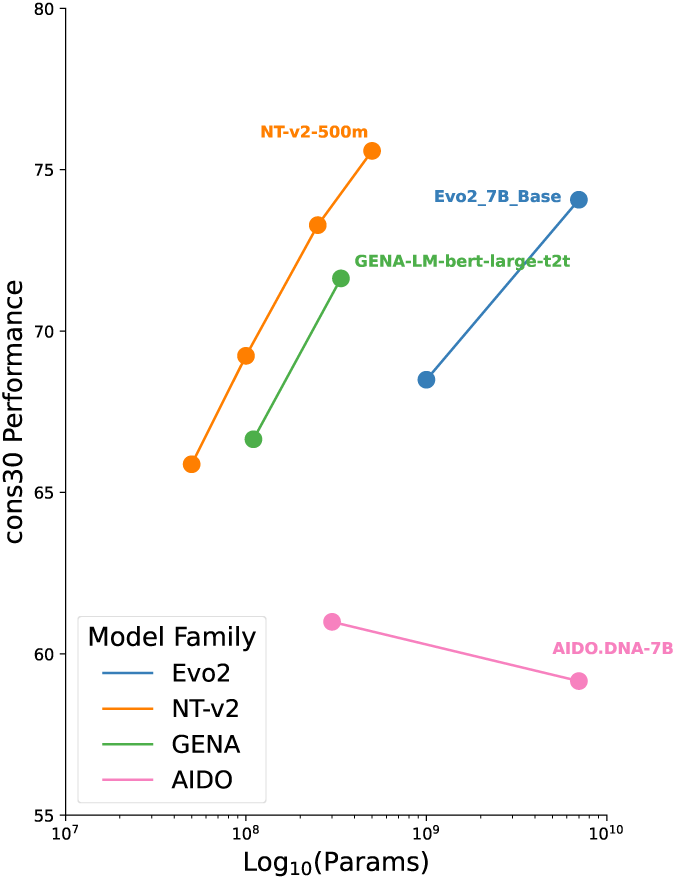
Scaling laws; scaling families. Conservation tasks demonstrate emergent *scaling laws* as a function of parameter count, when conditioned on the same training corpus & architecture. Different families, like Evo2 or GENA-LM, are less competitive than NT-v2, but scaling helps them offset this gap. AIDO models, on the other hand, diverge in both starting point and scaling results – we attribute this to the difficulty of training large-scale genomic models.

may have indicated a flawed training procedure as well. That is not to say that there is no value in these models, but `GUANinE` v1.1 is identify what that value may be.

### More context *̸*= better model

Next, we consider the trend of expanding input context [9, 10, 29, 30, 40]. For the tasks evaluated here, we note the performance of the 4 kbp-context Sei, which for functional annotation is nearly on par with the much larger 196 kbp-context Enformer and 524 kbp-context Borzoi models, see the `dnase-prop` and `ccre-prop` tasks in 2. We attribute Sei’s success to its parameter density, see “Digging into Sei” below for details.

Similarly, lightweight high-context architectures, such as the Mamba-style models in Caduceus, show promise, but we observe that the parameter-dense 500M version of Nucleotide Transformer v2 still achieves a higher performance on `cons30` and `cons100` at much lower context size, as shown in Table 2. At 65x the parameter count, NT-v2-500m features a *smaller* memory footprint with comparable FLoating-point OPerations (FLOPs) per example to Caduceus PS – making it a more readily trainable approach. Specifically, NT-v2-500m sits at 12.58 TFLOPs/example compared to Caduceus-PS at 7.84 TFLOPs/example [41]. On the same GPUs, we also managed to minibatch just 2 examples on Caduceus and 32 on NT-v2-500m.

In effect, increasing compute spent on context size is a tradeoff to spending it on parameter count or hyperparameter exploration.

### New task results

While our results suggest that the region-level conservation estimates of gLMs are currently more reliable than base-pair level annotations (see the strong performance of GPN-MSA and phyloP-30way on `cadd-snv` in Table 3), this strength of gLMs in conservation modeling extends to the level of individual variants as well. In this regard, we observe in Table 3 that neither Enformer nor Sei approach the performance of NT-v2-500m on `cadd-snv`. The performance of GPN-MSA on `cadd-snv` confirms that more information can be extracted from MSAs, which while not resolved by PhyloGPN, has nonetheless been suggested to improve sequence-only methods [23].

Turning to the ‘low’ performance of CADD v1.7, at least relative to two its inputs phyloP-30way and -100way, we note that per the CADD methodology [8, 34], the best-performing models on external validation data (not the training corpus) are only partially fit. Specifically, CADD 1.7 undergoes 13 iterations of L-BFGS and is stopped early to *prevent* convergence – far fewer iterations than standard. In var-CADD, this early stopping is even more pronounced, with its authors performing only ∼ 3 iterations of L-BFGS across training sets.

We highlight this intentional *under*fitting of CADD as the reason it outperforms the evaluated conservation-based approaches on `clinvar-snv`. We suspect this failure to generalize is due to both the intrinsic differences between deleteriousness and pathogenicity and to broader phenomena like the construct validity of ClinVar as a measure of pathogenicity [42]. In fact, while we do not report scores, we found that `cadd-snv` trained sequence models show little-to-no predictive power on `clinvar-snv` – a finding we did not anticipate originally.

### Evolutionary distal sequences can improve gLM performance

Next, we consider scaling the pretraining corpus, which is known to substantially impact the quality of the downstream model [38, 43, 44]. A reasonable question to ask for genomic language models is whether training on non-human sequences can improve human performance. We provide several points of evidence to suggest that yes, even distally related genomes, in sufficient quantity, can improve model performance on tasks of a different genome.

The most direct evidence comes from contrasting DNABERT-2 and Nucleotide Transformer v2 100M, as seen in Table 2. Given that the primary distinction between them is the much larger pretraining corpus of the latter, we attribute NT-v2-100M’s relative success over DNABERT-2 on to the additional gradients extracted from training on distally related genomes.

As a corollary, we observe that GENA-LM-bert-base-t2t-multi does not exhibit catastrophic forgetting on `GUANinE` tasks. GENA-LM-bert-base-t2t is fine-tuned down-stream on a combination of human and non-human genomes to produce the -multi model, yet the performance of the two is nearly equivalent.

Finally, we compare models trained on a small related genome versus using a much larger, unrelated genome for pretraining. Examining the `gpra-c` and `-d` task performance gap in Supplemental Table A1, we can see that between an hg38-trained model, GENA-LM-bert-base, and a yeast-genome trained model, GENA-LM-bert-base-yeast, the larger training dataset size of hg38 outperformed the more evolutionarily relevant yeast-trained model. Though a large collection of fungal genomes might alleviate this underperformance, this evidence suggests it is worth considering the size and number of distal genomes when pretraining gLMs.

### Digging into Evo2

Despite its superior performance at high-context sizes relative to other gLMs, Evo2 7B is one of the more difficult models to use. To assist other modelers, we detail a number of insights based on its performance on a large-scale benchmark like `GUANinE` .

Specifically, we found that Layer 26 of Evo2 7b performed the best across tasks. Embeddings extracted from layers 25 and 27 *rarely* exhibited better performance on the development set, and while we do not release test set results other from layers, we mention that this was usually due to overfitting, rather than truly better performance. More strikingly, we observed – across tasks – a strong decrease in performance using embeddings from deeper into the model. We believe this may be due to particular issues of its experimental architecture, be it missing residual connections or uniquely specialized later layers (output layers in LMs are known to be biased towards logits, rather than biological information [45]). We do not, however, observe anything near this performance degradation in any other model – including the similarly Mamba-based Caduceus models.

Additionally, we were able to infer single examples on 4x L40 GPUs^1^ at 32 kbp of input size by **deleting** deeper layers of the model, which we found to be necessary to achieve a reasonable runtime. We did not attempt to evaluate greater context sizes, despite the authors’ targeted 1 Mbp input size, due to computational infeasibility – we note that successful open 7B parameter architectures like Mistral [46] or Gemma2 [47] target more modest context sizes.

### Digging into Sei

Several unique factors drive Sei’s strong performance in spite of its small context size. We attribute its high parameter count (approaching 900M) and immense supervision – 21,907 output tracks – as the predominant factors in its performance.

Additionally, Sei is computationally lightweight despite its high parameter count – it requires 105.94 GFLOPs/example^2^ compared to Enformer’s single-bin inference of 31.16 TFLOPs/example. Note that Enformer parallelizes over genomic windows for training efficiency [9], which allows it to infer locus-centric *batches* much more efficiently^3^. Sei is also at a disadvantage when measuring FLOPs per bp – it achieves only 26.485 MFLOPS/bp compared to Enformer’s more rigorous 158.5 MFLOPS/bp.

However, this brings us to a key advantage of Sei: its supervision and model complexity. In fact, it tallies 222.5 Kparams/bp compared to Enformer’s 1.28 Kparams/bp [41]. We attribute Sei’s success to this increased model complexity, despite its lower input size and operational intensity [49].

### Combining supervised and unsupervised approaches for variant effect prediction

We also train and report results of a random forest (‘RF,’ `max depth=3`) ensemble of NT-v2-500m and Sei in Table 3. While it is possible to *linearly* combine the outputs of the two’s linear probes (ridge regression yields a test set score of 26.4942), we found it necessary to employ this non-linear approach to achieve more than marginal improvement over NT-v2-500m’s already strong results.

While properly combining supervised and unsupervised techniques is beyond the scope of this work, we suggest that such procedures likely need to be done *during* pretraining, although two-stage training as in GENA-LM-t2t-bert-base-enformer may be simplest.

## Discussion

No one model can accomplish every task, and our results highlight the need for large-scale benchmarks like `GUANinE` , unhindered by sample size concerns or reporting bias, to evaluate pretrained genomic models across a variety of tasks. Our results also reveal general trends in performance with respect to modeling choices such as input context size and parameter count.

Controlling for other variables, context size should improve model performance; however, we note that models like Evo2 7B achieve a Pyrrhic victory in this regard – research groups attempting to use this model will likely be restricted to either a small sample size or a shallow analysis, given the demanding compute requirements for even a single inference. Specifically, in the evaluation of `GUANinE` v1.1, we utilized a total of approximately 1.9 zetta-FLOPs of compute – 80% of which was dedicated to Evo2 models alone.

Individual models effectively have very different use cases. Consider AlphaGenome, which for a clinical geneticist could reasonably infer a few thousand variant effects, or Evo2, which for a synthetic biologist could embed tens of thousands of synthetic sequences – neither is ideal for an academic computationalist hoping to experiment on model behavior through a few millions of inferences. For this latter use case, we would point modelers to NT-v2 or Sei instead.

In particular, we note the combined utility of Sei-Aggr, which is simplifed to the near-zero-shot regime through the variant scoring approach presented by Avsec et al. [10] (see Methods for our implementation) and Pangolin – the two together see 10 kbp of context, but outperform AlphaGenome’s 1 Mbp of context on `clinvar-snv`. This finding is in line with previous reports that large-context models do not fully make use of distal sequence context [13].

Our results showing complementary strengths of supervised models and language models on different tasks suggest that combining these modeling strategies may be a promising approach. A predominant paradigm in the field of natural language processing^4^ is the self-supervised pretraining of a large model, which can then be supervised and fine-tuned on downstream tasks [37, 43]. This was preliminarily investigated on a task-by-task basis in `GUANinE` v1.0 through the introduction of the hgT5 models [3].

With the larger set of `GUANinE` v1.1 evaluations, we can now point to the success of GENA-LM-bert-base-t2t-enformer, which was fine-tuned by its developers on the same training data as Enformer. As a result, this downstream model outperforms its original on both functional element annotation and conserved sequence modeling, although given its slight decrease in performance on `gpra-c` and `-d`, it appears to gain specialization to human sequences as a result.

We further demonstrate a proof-of-principle hybrid-supervised approach in Table 3, where a random forest ensemble of NT-v2-500m and Sei was able to outperform either alone. While this result confirms that each has something to learn from the other, the modest gains from this initial ensembling approach suggest hybrid models may be difficult to implement in practice. We are nonetheless optimistic that future work can validate such approaches.

## Conclusion

The version 1.1 expansion of the `GUANinE` genomic sequence-to-function benchmark has illuminated dozens of novel insights for the field of genome sequence modeling. Chief among them is clarity regarding the roles of supervised sequence-to-function and unsupervised genomic language models: the two have distinct strengths and use cases, and since both allow for unique perspectives on variant effects, we believe the two can be intertwined in the field going forward to enable models to more fully comprehend a genomic sequence or the effects of variation.

## Methods

### Evaluated models

#### ChromBPNet models

Pampari et al. [22] introduced compact, cell-line-specific supervised models tailored to *decision explainability*, i.e. identifying putatively causal motifs. The ChromBPnet family of models are bp-resolution CNNs that accept 2,114 bp of input and output a factorized 1,000 bp of signal. We choose to include two folds of the ATAC-seq trained models based on the GM12878 cell line, which has a high 572M read depth. Fold 0 is the preferred model in Pampari et al. [22], although we also report the *unbiased* Fold 3, which is consistent with `GUANinE` ’s train-test split. As the scalar output is less transferable to tasks in `GUANinE` v1.0, we pruned layers after `add 7` and used the ultimate 512-dimensional embeddings for ChromBPNet models. This leaves a total of ≈ 6.34M parameters, much smaller than other models.

#### ENCFF180XQC baseline

In addition to the ChromBPNet models, we evaluate the pre-processed signal input to the models, namely, ENCODE track ENCFF180XQC. We report this baseline to give readers insight into the ATAC-seq chromatin accessibility signals underpinning `GUANinE` tasks (i.e. construct validity), as well as to provide a new form of baseline – raw experimental data.

#### DNABERT-2

A followup to the early gLM DNABERT, the second version maintains a comparable performance footprint but at higher context sizes through byte-pair encoding tokenization. The model is trained via unsupervised masked language modeling across 135 species spanning multiple phylogenetic kingdoms – without any apparent deduplication (the authors merely remove sequences with ‘N’). DNABERT-2 follows the original BERT-base architecture but with minor tweaks, such as a position-aware attention mechanism. We aggregate per-bp final-layer embeddings of dimension 768, and the model consists of 117M parameters.

#### Nucleotide Transformers v2

Trained across a broad set of 850 species, the second iteration of the Nucleotide Transformer gLMs use k-mer tokens (predominantly 6-mers) on moderate context sizes with an enhanced BERT-style architecture. Trained via masked language modeling, the corpus of NT models is an augmented with observed variation, including individual haplotypes from 1000 Genomes [53]. Similar to DNABERT-2, there is otherwise minimal filtering of the training corpus. Model parameters in the v2 family are included in the name of each model, e.g. ‘100M,’ and correspond to hidden dimensions of 512, 512, 768, and 1024, respectively. We abbreviate these models ’NT-v2’ throughout this work.

#### ESA-dna2vec

This contrastive model is distinct for its dedication to sequence embedding – and its usage of only ≈ 2% of hg38 during training [26]. While it is relatively underpowered compared to other methods, it showcases the potential for contrastive pretraining in genome modeling. The model sits at a total of ≈ 54M parameters, of which the initial embedding layer constitutes 18.9%.

#### PhyloGPN

This MSA-inspired model is specialized to predict centered SNVs given 480 bp of surrounding context. While the training loss is weighted via MSA information, forward inference is performed on sequences alone, enabling its straightforward evaluation as a sequence model in `GUANinE` . The model totals just over 83 million parameters, of which the output logit layers make up ≈ 0.007%. Aside from the `clinvar-snv` task, which logits are ideal for, we use the 960-dimension mean-pooled embeddings. We report both embedding and zero-shot logit scores for `cadd-snv`.

#### GENA-LM

A family of models and architectures, the GENA-LM series uses byte-pair encoding across different genomes, including the human t2t genome [54], during gLM pretraining. Pretraining genomes were typically augmented with SNVs from 1000 Genomes or similar. Architectures span BERT-base, BERT-large, and BigBird, which offers a much higher input context than traditional transformer encoders. We do not evaluate the sparse implementation of BigBird, but those willing to install it will likely see speedups. Of architectures evaluated, the models consist of 110M, 336M, and 110M parameters, respectively.

##### Caduceus

The two Caduceus gLMs evaluated, PH and PS, evaluate reverse-complement-augmented and RC-equivariant architectures, respectively, using versions of the long-range Mamba SSM architecture [29, 55]. Both models are trained with only the hg38 reference genome – a significant disadvantage compared to many other gLMs in `GUANinE` v1.1. Each model consists of ≈ 7.7M parameters, making them among of the smallest high-context models evaluated.

##### Evo2

A slow yet potent family of gLMs evaluated, the Evo2 models use a custom SSM-esque architecture to achieve high context size inference on a broad, yet pruned, set of genomes. We do not attempt to infer Evo2 7B beyond 32 kbp of input, nor do we evaluate Evo2 40B, as for a *single example*, either requires more than 192 GB of VRAM on Lovelace+ Nvidia GPUs (computational constraints limited us to 4x L40 GPUs). The 1b base and 7B base models operate on 8.2 kbp input sizes; we found that uncased (uppercase) sequences yielded much better performance despite the models having been trained on repeat-masked sequences.

##### Sei

The third iteration of DeepSEA, Sei continues the tradition of flattening a large convolutional network into an output layer – albeit with a more sophisticated non-linear output than previous versions. Uniquely, Sei also predicts 21,907 tracks, making it the most-supervised model evaluated in v1.1, although Borzoi or AlphaGenome have different types of data compared to Sei (e.g. RNA-seq or splicing, while Sei is primarily ChIP-seq). Being intensely concentrated on its 4 kbp input size, Sei consists of nearly 890M parameters, of which 91.7% are located in the quadratic output layers. Given this density of weights in the output layers, we use outputs, not embeddings, for all Sei results.

#### AIDO

As part of a larger AI Digital Organism (‘AIDO’) project for model generation, AIDO.DNA models are gLMs trained on nearly the same corpus as the Nucleotide Transformer models. Both the 300M and 7B formulations are trained on 4 kbp of input. In contrast to the reported results of Ellington et al. [31], we do not fine-tune the full models, nor do we select the optimal checkpoint for each evaluation task. This may partially explain the near-equivalence of the two’s performance in our results.

#### Mistral-DNA and ModernBERT

Pedagogically-inclined models developed by Raphael Mourad [28], Mistral-DNA and ModernBERT use sophisticated architectures on minimally preprocessed hg38 datasets, but still achieve respectable scores on `GUANinE` tasks. We attribute this not solely to the power of modern LM architectures, but also to the underexplored hyperparameter space of gLMs more generally.

#### GPN-MSA

A multiple-sequence-alignment-to-function model, GPN-MSA is a sophisticated approach to sequence conservation modeling. Due to its not directly modeling un-aligned sequences, we do not attempt to evaluate GPN-MSA on most tasks – instead, we use the precomputed genome-wide scores computed by Benegas et al. [33]. See https://huggingface.co/datasets/songlab/gpn-msa-hg38-scores.

### Datasets

#### clinvar-snv

The `clinvar-snv` dataset was designed to be harmonious with existing pathogenicity and deleteriousness detection methods, rather than to be a (possibly ill-defined) objective measure of pathogenicity grading.

To that end, we began by cross-referencing the 2+ star non-coding variants used in CADD v1.7 (compiled June 2023, [8]) with the more up-to-date set of ClinVar variants from 24 August of 2025. Our test set consists of only the pathogenic and benign variants present in both (i.e. still considered accurate), while our development set is are those added to ClinVar since June 2023 – and thus not originally evaluated in CADD v1.7. This guarantees that CADD v1.7, and models fairly compared on our test set, do not benefit from data leakage when reporting scores on our `clinvar-snv` task.

This yields an imbalanced set of 58899 dev benign, 4601 dev pathogenic, 28900 test benign, and 5982 test pathogenic 2+ star variants, which we sought to class-balance via downsampling. Given the extreme feature imbalances present in ClinVar (substitution rate, VEP label, chromosome), we implement propensity score matching [56] across the aforementioned features to downsample an optimized, class-balanced subset of benign variants. Propensity models were an ensemble of seven randomly sampled, class-balanced logistic regressions. Subsets were then derived by matching propensity scores via PuLP [57] and set to the maximum possible balanced number of pathogenic variants per chromosome.

#### cadd-snv

We begin with the substitution-rate-balanced hfs set of VarCadd [34], available in the VarCADD Zenodo repository (link). This corresponds to a 50/50 split of proxy benign SNVs and proxy deleterious singletons (`s`), the former being a combination of evolutionary-derived variants (`h`) and frequent variants (`h`).

Four post-processing filters were then applied:

1. Removing coding variants (located in ‘CDS’ regions per GENCODE v48 [58])
2. Deleting mislabeled *variants* according to ClinVar (e.g. removing CADD’s proxy benign variants listed in ClinVar’s ‘Pathogenic’ set)
3. Pruning leaked *positions* in ClinVar (e.g. coordinates present in the proxy benign set with a benign ClinVar alt-allele)
4. Downsampling proxy benign and proxy deleterious variants to be balanced per chromosome-substitution (e.g. ‘Chr 22 G*>*A’).

These specific filters imply a number of unique properties about the resulting data, most notably the inclusion of splicing variants and those located in UTR regions. Additionally, the orthogonalization of ClinVar and `cadd-snv` variants ensures that models trained or validated using `cadd-snv` have reduced risk of overfitting to `clinvar-snv`. We finally note that our chromosome-substitution balancing is more rigorous than the original VarCADD procedure, although this represents an additional ≈ 2% downsampling for `cadd-snv`.

While a larger yet still balanced dataset could be achieved via filtering the `hfs-all` dataset, we opine that `GUANinE` is meant to be a benchmark, not a comprehensive training database. We would encourage model developers seeking to maximize performance to assemble the highest quality training data, then screen it against `GUANinE` ’s dev and test splits for comparison.

Dataset reductions, in order of derived, frequent, and (deleterious) singleton sets, are roughly:

1. Prune coding variants: ≈ −0.6%, −0.8%, −1.2%
2. Filter against any-star ClinVar: -95, -108, -2646 (all less than 0.01%)
3. Downsampling to rebalance: ≈ −2.2%, −2.2%, −1.8%

#### Few-shot cadd-snv

As in the original `GUANinE` benchmarks, we derive a ‘few-shot’ subset of the training data which corresponds to ≈ 1% of the original training size [3, 59]. To do this, we use perform stratified sampling proportionately on chromosome-substitution combinations to preserve, across features, the class balance of proxy benign vs proxy deleterious. In terms of bias-variance tradeoff, this reduces the variability of models fit to the few-shot subset, which should more accurately resemble said model’s performance on the larger training set. Modelers should note this further entrenches the *per-observation* biases of the `cadd-snv` dataset^5^ – those who are seeking to predict deleteriousness across *samples* would benefit from alternative sampling procedures (i.e. different frequencies of sampled variants).

### Evaluation methods

The key to successful linear evaluation is selecting the proper regularization strength for the Ridge regression. In `GUANinE` , we sweep over a wide range of values during 1% ‘training,’ spanning *C* ∈ [0.01, 0.03, 0.1, · · · , 3000., 1e5, 3e5] for Ridge regression [60] to select the best performance on the development set. For logistic regression, which uses an inverted *α*, we used *α* ∈ [1e-6, 3e-6, 1e-5, · · · , 0.1, 0.3, 1]. The best performing model on the development set was selected for test set inference.

Due to its high dimensionality, Sei uniquely required additional regularization, leading us to expand this hyperparameter search to the above ranges. The same occured with the ridge-ensemble of NT-v2-500M and Sei, which while not material, had a large value of *α* = 1e13.

For the `cadd-snv` task, we use the log2 fold change difference of sigmoid-functioned outputs (e.g. Sei) when possible. Alternatively, especially for gLM embeddings, we simply use the alt - ref contrast. Other than for zero-shot performances of phyloGPN and GPN-MSA, we do not use gLM output logits.

Finally, we use last-layer embeddings for all gLMs (except Evo2, as noted in the Discussion), and primarily use outputs for supervised models, when possible. Notable supervised exceptions include the ‘pre-output’ Enformer embeddings and both the ChromBPNet models given the low generalizability of its outputs. We suspect that like Enformer, many supervised models may have better performance at penultimate layers, but we do not believe this to be the case for DeepSEA, Beluga, or Sei, given their top-heaviness.

### Calibrating AlphaGenome and Sei-Aggr

As described, but not detailed, in the AlphaGenome preprint [10], we utilized a small-N ‘calibration’ of combined variant scorers derived by aggregating the output of models by experiment type. For Sei, we found that aggregating experiments with at least 20-50 examples, coupled with pruning the rarer experiments, produced the best results (using ‘20’ as the threshold shrinks the dimensionality from 21,907 → 109).

For AlphaGenome, we selected all tracks for a single cell line for aggregation, the well-represented K562 line, per the guidance by the uathors[10].

Once a low-dimensional scorer is achieved, for either the K562-outputs of AlphaGenome or Sei, we finish calibration by using logistic regression fit to the development set to weight each experiment type into a final score, similar to Avsec et al. [10].

### Logarithmic feature intervals

As has been suggested by model developers [9, 40] and evaluators [61], inferencing with alternative-resolution-pooled features is essential for thorough model evaluation. For new models in `GUANinE` v1.1, we utilized a sequence of concentric, single bp-centered resolutions for embeddings or outputs (where possible, e.g. not Sei or variant approaches like AlphaGenome). These correspond to a sequence of *L*(*k*) = 1 + 2*^k^* bp, or a series of [1, 3, 5, 9, 17, 33, 65, · · · , ⌈log_2_Context Size⌉] regions, with the final being equivalent to mean-pooling.

Note that for variable token-length models like DNABERT-2 or the NT-v2 family, we use partially-weighted tokens when region boundaries fall in the middle of a token (e.g. 1 bp of a 3bp token = 1*/*3).

For coarse resolution models like Enformer, we report the best of 1 or 3 bins, etc, see previous work for details [61].

### Notes on Spearman’s *ρ*

We unconventionally choose to report Spearman’s *ρ* for our Boolean classification tasks, cadd-snv and clinvar-snv . We justify this with both the ease of interpretability (as *ρ*^2^ is the proportion of variance explained by ranks), and the fact that it reduces to the point-biserial correlation, a well-studied distribution for class-balanced situations such as ours [62]. Additionally, the point-biserial correlation is known to be monotonically related to AUROC [63], with the latter being more common but less *directly* interpretable. We point readers interested in the numerical comparison of metrics to Salgado [63].

Wherever possible, we use continuous outputs for scoring, i.e. `predict_proba` for scikit-learn models like logistic regression. This relaxed predictor provides both better performance and more accurately captures a model’s estimates.

## Availability of data and materials

The `GUANinE` benchmark can be accessed on HuggingFace at https://huggingface.co/ `GUANinE`. User guides, explainers, and instructions can be found on the semi project website at https://guanine.readthedocs.io/en/latest/.

## Acknowledgements

Special thanks to Ian Holmes for editorial feedback. Thanks to Ayesha Bajwa and other members of the Ioannidis group for insights on model results.

## Ethics Declarations

### Ethics approval and consent to participate

Not applicable.

### Consent for publication

Not applicable.

## Appendix A Extended results

**Table A1:**
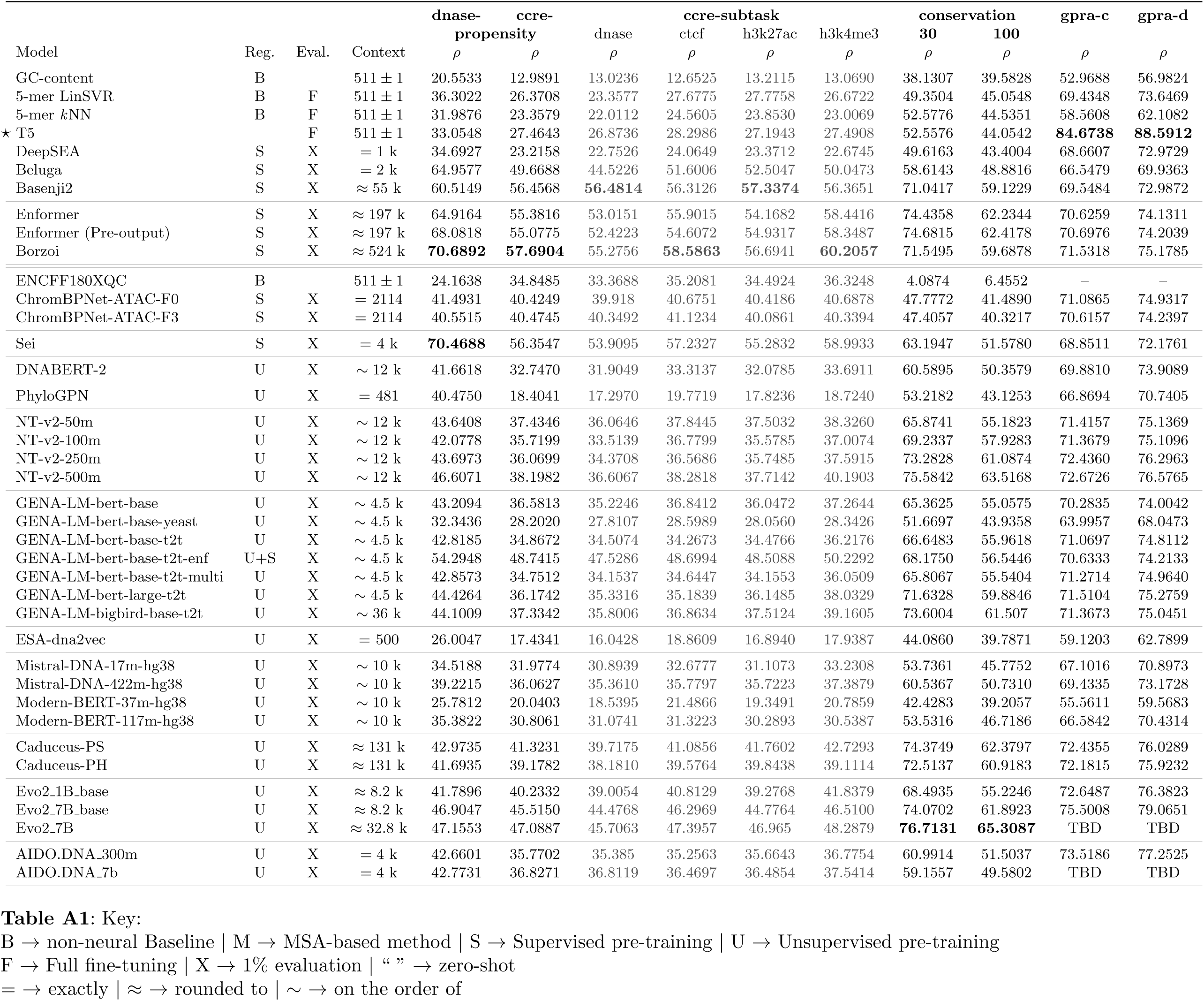
Key: B → non-neural Baseline | M → MSA-based method | S → Supervised pre-training | U → Unsupervised pre-training F → Full fine-tuning | X → 1% evaluation | “ ” → zero-shot = → exactly | ≈ → rounded to | ∼ → on the order of

## Appendix B Supplemental Background

Here we detail summaries of the six different tasks from `GUANinE` v1.0, grouped by type.

1. Regulatory elements: dnase-prop asks a model to assess tissue-agnostic DNase-hypersensitivity, corresponding to ENCODE v3’s definition of a DNase-hypersensitive site [19]. Input sequences are taken from hg38, with the output being an ordinal 0-4 scalar of consensus-peak accessibility. Its corollary, `ccre-prop` instead requires estimating the *functional properties* of those accessible sequences, such as candidate cis-regulatory element (cCRE) histone modifications like H3K4me3 [19, 64]). Many supervised models, from DeepSEA to Enformer, have seen the training and even testing data for these two tasks, making them in-distribution [61].

2. Conservation: `cons30` and `-100` require estimating the degree of conservation, or tendency to stasis, of an input hg38 sequence using an aggregated score of the popular phyloP metric [20]. The 30 consists of conservation defined across a 30-way mammalian multiple sequence alignment, primarily including primates, while the 100 consists of a 100-way alignment of vertebrate sequences, primarily mammals.

3. Synthetic promoters: The two synthetic tasks, `gpra-c` (complex) and `-d` (defined) correspond to two sets of large scale dual-reporter assay experiments measuring *gene expression* [21, 65]. Randomized promoters are prepended to reporter genes before being transformed into yeast.

Continuing in the direction of benchmark longevity, most tasks in `GUANinE` do not have publicly accessible test *labels*, given the risks of (inadvertently) overfitting to test sets, as well-documented in Kaggle contests [66].

Let us also enumerate previously reported model performances, which list the paper detailing their evaluation (see respective manuscripts for model details):

1. GC-content, 5-mer Linear SVR, 5-mer kNN non-neural baselines [3]
2. T5 and hgT5 neural baselines [3]
3. DeepSEA, Beluga, and Basenji2 supervised CNNs [3]
4. Enformer and Borzoi supervised hybrid-convolutional models [61]

Note that as `cadd-snv` and `clinvar-snv` have not previously been described, we novelly report results on those tasks for several of these models.

## Footnotes

1 Specific architectural choices prevent Evo2 models from being inferenced on GPU architectures released by Nvidia before 2022.

2 We use the standard M = *Mega*, G = *giga*, and T = *tera* prefixes when tallying FLoating-point OPerations (FLOPs [48]).

3 For instance, inferring three consecutive bins is a marginal increase to 32.2 TFLOPS/example – this trend holds to approximately .02 TFLOPS per additional bin [41].

4 ^4^This trend began in the supervised regime of computer vision [50–52], but due to the modern dominance of masked language modeling, is better exemplified in NLP.

5 ^5^Deleteriousness is not evenly distributed across substitutions, which themselves are variable across genomes.

